# Disruption of *Histone H4C* genes impairs skeletal development and cortical neurogenesis, modeling rare neurodevelopmental syndromes

**DOI:** 10.64898/2026.07.12.738071

**Authors:** Haruna Nagasawa, Kaneyasu Nishimura, Sayaka Tojima, Tadashi Nomura

**Affiliations:** Applied Biology, Kyoto Institute of Technology, 1 Hashikami-cho, Matsugasaki, Sakyo-ku, Kyoto, 6068585, Japan; Laboratory of Functional Brain Circuit Construction, Graduate School of Brain Science, Doshisha University, 1-3 Tatara Miyakodani, Kyotanabe, Kyoto, 6100394, Japan; Center for Social and Biomedical Engineering, Kyoto Institute of Technology, 1 Hashikami-cho, Matsugasaki, Sakyo-ku, Kyoto, 6068585, Japan

**Keywords:** Histone H4, Neural progenitors, Cortical organoid, Genome-editing, TEBIVANED Syndrome

## Abstract

Histone proteins, which reside in the nuclei of eukaryotic cells, are involved in diverse cellular processes. The core histone H4 serves as a structural component of the nucleosome. Patients carrying mutations in H4Clustered histone (*H4C*) genes exhibit a broad spectrum of developmental abnormalities, including short stature, microcephaly, intellectual disability, growth retardation, and digital anomalies. However, the impact of H4 mutations on mammalian embryogenesis remains largely unclear. Here, we demonstrate that *histone H4C* genes play crucial roles in skeletal development and cortical neurogenesis. We found that mRNAs of the *histone H4C* gene family are specifically expressed in proliferating progenitor cells in the developing mouse neocortex and in human induced pluripotent stem cell-derived cortical organoids. CRISPR-mediated disruption of *H4C3* in mice caused severe defects in skeletal formation and neocortical neurogenesis. Furthermore, overexpression of a mutant form of *H4C3* resulted in altered expression of genes associated with cellular migration and motility. Together, these findings suggest that histone H4 plays a critical role in regulating the balance between proliferation and differentiation during mammalian embryonic development, thereby explaining the broad spectrum of patient phenotypes.

## 1. Introduction

The DNA present within the nucleus of eukaryotic cells is packaged into a chromatin structure. Nucleosomes, which constitute chromatin, are repetitive structures consisting of double-stranded DNA wrapped around histone proteins (Bannister and Kouzarides, 2011; Kornberg, 1974; Luger et al., 1997). After translation, histone proteins undergo various modifications, such as acetylation, phosphorylation, and methylation. These chemical modifications influence the interaction between histone proteins and DNA, thereby altering the structure of chromatin (Allfrey et al., 1964). The structure and function of these histone proteins play a crucial role in individual development and maturation by regulating cell-specific processes such as DNA replication, transcription, and repair (Malone et al., 2026).

There are five types of histone proteins: H1, H2A, H2B, H3, and H4. H1 is called a linker histone and binds to the linker DNA region that connects nucleosomes (Baxevanis and Landsman, 1996). H2A, H2B, H3, and H4 are called core histones; they form octamers and function as the structural units of nucleosomes. The amino acid sequence of H1 is highly conserved in the globular domains but is generally less conserved overall. In contrast, the amino acid sequences of core histones are highly conserved, with H4 exhibiting particularly high conservation.

Human histone H4 is encoded by 15 histone H4 genes, while mouse histone H4 is encoded by 13 histone H4 genes (Seal et al., 2022). Human histone H4 genes are distributed across three chromosomes. All histone H4 genes encode the same amino acid sequence. The amino acid sequence of the human histone H4 gene is highly conserved across the animal kingdom and is 100% conserved between humans and mice (Baxevanis and Landsman, 1996). This suggests that there are strong evolutionary constraints on the structure and function of the histone H4 protein.

Recently, congenital disorders caused by a single amino acid substitution in human *H4C* genes have been reported (Tessadori et al., 2022; Tessadori et al., 2017; Tessadori et al., 2020; Tudorache et al., 2026). Patients with these mutations exhibit various abnormalities such as short stature, microcephaly, intellectual disability, growth delay, and foot abnormalities. Overexpression of a mutant H4 (H4K91Q) into zebrafish embryos results in severe morphological abnormalities, partially recapitulating symptoms observed in humans patients (Tessadori et al., 2017; Tessadori et al., 2020). Furthermore, a recent study reported that H4K91 mutant protein enhanced chromatin accessibility and altered developmental gene expression, which result in precocious neuronal differentiation (Feng et al., 2023). However, since the expression patterns of the Histone H4C genes has not yet been fully characterized, it remains largely unclear how abnormalities in these genes influence brain and body morphogenesis.

Here, we report that histone *H4C* genes play crucial roles in mouse embryogenesis. We demonstrate that H4C genes are specifically transcribed in proliferating progenitor cells in the developing mouse neocortex and in human induced pluripotent stem (iPS) cell-derived cortical organoids. CRISPR-mediated disruption of H4C3 in mice caused severe defects in skeletal formation and neocortical neurogenesis. Furthermore, H4K91Q, a mutant form of H4C3 altered expression of genes associated with cell adhesion, which were corroborated by *in utero* misexpression of the mutant H4C. Our findings suggest that histone H4 plays a critical role in regulating the balance between proliferation and differentiation during embryogenesis, thereby explaining the broad spectrum of patient phenotypes.

## 2. Methods

### 2.1 Animals

Pregnant female mice (CD-1 background) that were originally obtained from Japan SLC were maintained in a 12 h dark/light cycle at Kyoto Institute of Technology. Noon of the day the vaginal plug was identified was designated as E0.5.

### 2.2 Plasmid construction

Full-length cDNAs of mouse and human *H4C* genes were obtained from mouse genome by PCR-based amplification. Mouse *Tbr2* (*Eomes*) was subcloned from pCAG-Tbr2 vector (Nomura et al., 2016). Wild-type and a mutant form (K91Q) of mouse H4C3 were obtained by gene synthesis (gBlocks, Integrated DNA Technologies) and subcloned into the pBluescript SK or pCAG-RB vector by using an In-Fusion HD cloning kit (Takara Bio). The sequences of PCR primers were described in Table S1.

### 2.3 Labeling of S-phase cells

To label S-phase cells, Bromodeoxyuridine (BrdU) dissolved in PBS (10 mg/mL) was administered intraperitoneally at a dose of 0.5 mL to pregnant mice 1 hour before fixation.

### 2.4 Immunohistochemistry

Mouse embryos and human iPS cell-derived organoids were fixed with 4% paraformaldehyde dissolved in phosphate-buffered saline (PBS) at 4°C overnight for mouse embryos and 30 min for human iPS cell-derived organoids. After PBS washes, the embryos were cryoprotected with a 20% sucrose solution and immersed in Tissue-Tek. The frozen samples were sectioned at a thickness of 20 µm using a cryostat (Leica CM1850, Germany), and incubated with primary antibodies, including anti-Pax6 (rabbit polyclonal, PD022, MBL), Histone H4 (rabbit polyclonal, AB177840, abcam), βIII-tubulin (mouse monoclonal, AB9354, Merck Millipore), GFP (rat monoclonal, 04404-26, Nacalai tesque) and BrdU (mouse monoclonal, B35128, Invitrogen) antibodies. To detect BrdU, the sections were incubated with 2N HCl at 37°C for 15 minutes. After washing, the sections were incubated with secondary antibodies, including Alexa-Fluor 488 or 555 conjugated anti-rat or anti-rabbit antibodies (A11006, A21428, A21235, Invitrogen).

### 2.4 *In situ* hybridization

Whole mount and section *in situ* hybridization was performed according to a previous report (Osumi et al., 1997). After the amplification of the *H4C* fragment containing T7 and T3 promoter region with M13 primer, sense and antisense RNA probes were made by using T7 or T3 RNA polymerase (ROCHE). T3 or T7 polymerase were used for making digoxygenin (DIG)-labeled sense and antisense RNA probes. The samples were treated with proteinase K (10 min, room temperature, QIAGEN) and hybridized with riboprobe in hybridization buffer (50% formamide, 20×SSC, 10% SDS) at 68℃ overnight. Section slides were washed with Phosphate-Buffered Saline + 0.1% tween 20 solution (PBT) and diethylpyrocarbonate (DEPC)-treated deionized water. The samples were washed 3 times with 5×SSC and 2×SSC (50% formamide, 20×SSC, 10% SDS) and incubated with anti-DIG-AP Fab fragments (Merck) at 4℃ overnight. All the sections were visualized by NBT (Nitroblue Tetrazolium, Merck) and BCIP (5-Bromo-4-chloro-3-indolyl-phosphate, Merck) reaction.

### 2.5 Generation of cortical organoids derived from human iPS cells

The maintenance of human iPS cells and the generation of organoids were performed at Doshisha University using a modified version of a previously reported method (Amimoto et al., 2021). Human iPS cells [hiPSC line 1231A3; (Nakagawa et al., 2014)] were obtained from Center for iPS Cell Research (Kyoto University) via the RIKEN BioResource Center. The iPS cells were cultured in Essential 8 medium (Thermo Fisher Scientific) supplemented with 1% penicillin-streptomycin (Fujifilm Wako Pure Chemical Corporation) in culture dishes coated with iMatrix-511 Silk (Nippi, Tokyo, Japan). For passaging, cells were seeded at a density of 1,000–2,000 cells/cm² in Essential 8 medium supplemented with 1% penicillin-streptomycin. 10 μM Y-27632 (Selleck Chemicals) was added the culture medium for the first 24 hours after replating.

Cortical organoids were generated in three-dimensional culture using the SFEBq method (Eiraku et al., 2008). In brief, human iPS cells were dissociated using 0.5×TrypLE Select (Thermo Fisher Scientific) and seeded onto U-bottom 96-well plates (Sumitomo Bakelite, MS-9096U) in the differentiation medium described below (9,000 cells per well, 200 μL) to form cell aggregates.

Subsequently, the cells were differentiated in Essential 6 medium (Thermo Fisher Scientific) containing 1% GlutaMax (Thermo Fisher Scientific), 1% non-essential amino acids (Fujifilm Wako Pure Chemical Corporation), 0.1 mM 2-mercaptoethanol (Fujifilm Wako Pure Chemical Corporation) and 1% penicillin-streptomycin. To this Essential 6 medium, 200 nM LDN193189 (days 0-11, Stemgent), 500 nM A83-01 (days 0-6, Fujifilm Wako Pure Chemical Corporation), 2 µM XAV939 (days 0-11, Selleck Chemicals), and 10 µM Y-27632 were added starting on day 0, and Y-27632 was removed from the medium on day 3. From day 5 to day 11, the medium was gradually replaced from Essential 6 medium to AscleStem Neural Medium (Nacalai Tesque) supplemented with AscleStem Neuronal Supplement (Nacalai Tesque), 1% GlutaMax and 1% penicillin-streptomycin.

From day 11, 20 ng/mL brain-derived neurotrophic factor (BDNF, PeproTech) and 200 μM ascorbic acid (Sigma-Aldrich) were added. Half of the medium was replaced every other day, and induction was continued until day 76.

### 2.6 Mouse genome engineering

Genome edited mouse with mutated H4C3 alleles were established by CRISPR/Cas12a(Cpf1)-mediated homology directed repair (HDR). A guide RNA sequence that targets mouse H4C3 was designed by CHOPCHOP (https://chopchop.cbu.uib.no). We chose a target sequence with no off targets based on mouse genome information (mm10/GRCm38). As an HDR template for homologous recombination, single strand DNA (ssDNA) attached with Myc-tag sequences was designed.

Genome-edited mice on the CD-1 background were generated by the *i*-GONAD method (Ohtsuka et al., 2018). Briefly, pregnant female mice (E0.75) were anesthetized with 2% isoflurane and the CRISPR/Cas12a solution was injected into the oviduct lumen. Then, square electric pulses were applied to the oviduct by using a pulse generator (super electroporator NEPA21, Nepa gene). All sequences of guide RNAs and PCR primers were depicted in Table S1.

### 2.7 Skeletal analyses

Embryonic mice were collected and dissected in phosphate buffered saline. The samples were fixed with 100% ethanol for several days and degreased by acetone overnight. Skeletal staining was performed by Alcian blue and Alizarin red according to standard protocols (Rigueur and Lyons, 2014; Saito et al., 2025).

### 2.8 RNA-seq analyses

Mouse neuronal cells were isolated from E13.5 embryonic neocortex. After introduction of the expression vector (pCAG-H4C3WT or pCAG-H4C3K91Q) by electroporation with a cuvette electrode (SE-201, BEX), the cells were cultured in Neurobasal medium (Thermo Fisher Scientific) supplemented with GlutaMax, B27, and human FGF-2 (Thermo Fisher Scientific) at 37 °C, 5% CO_2_ for 24 hours. After the culture, total RNA was prepared from the cells by using FastGene RNA premium kit. Residual DNA was eliminated by DNase treatment. The quality and quantity of RNAs were assessed by using an Agilent Technologies 2100 Bioanalyzer or 2200 TapeStation (Agilent). The cDNA library was constructed using a TrueSeq Standard mRNA LT Sample Prep Kit according to the manufacturer’s protocol (15031047 Rev. E) and sequenced on an Illumina platform and 101 bp paired-end reads were generated. The sequence data were mapped to a reference genome sequence (*Mus musculus* GRCm38, GCA_000001635.2) with a splice-aware aligner (HISAT2 v 2.1.0). The transcripts were assembled by StringTie (v 2.1.3b) with aligned reads. GO enrichment analysis was performed based on Gene Ontology.

### 2.9 Analysis by using public database

To examine the expression pattern of *H4C3* (*HISTH1H4C*) in human embryo, we used a public database [Spatiotemporal transcriptome atlas of the developing human brain (http://donglab.life/brainAtlas.html#target-div)]. To comprehensively analyze the expression of histone H4C genes in the mouse neocortex, we utilized the dataset from 10xGENOMICS (10k Mouse E18 Combined Cortex, Hippocampus and Subventricular Zone Cells, Chromium GEM-X Single Cell 3’) and compared the expression of histone gene clusters and cell-specific marker genes across each cluster by using Loupe Browser 8.1.0. Published transcriptome data of neural cells differentiated from mouse ES cells (Feng et al., 2023) were used for re-analysis of genes affected by H4C3K91Q. Generally applicable gene-enrichment analysis (GAGE) was conducted by iDEP and iDEP2.

### 2.10 Image processing and statistical analysis

Sample images were captured by using the microscope, (TF-B, Kennis; SZX7, Olympus; ECLIPSE Ni, Nikon; M165 FC, LEICA or FLUOVIEW FV4000, Olympus) equipped with a cooled CCD camera (DEGITAL SIGHT DS-Fi2, Nikon, DsFi1c, Nikon or DS-F/3, Nikon). All captured images were processed with Image J (1.53k) and Photoshop (v2025, Adobe), CellSens FV (ver3.1.1.67). For statistical analysis, the results of three independent experiments were compared. To quantify GFP fluorescent intensities at the endofeets, background intensity was subtracted from the signal intensity.

Comparisons between experimental groups were performed using Microsoft Excel (v16.98, Microsoft) and Prism 9 (v10.5.0, GraphPad9). Statistical significance was determined using unpaired Welch’s t-test and 2-way ANOVA with Šídák’s multiple comparisons test.

## 3. Results

### 3.1 Cell type-specific expression of *H4C* genes in the developing mouse and human neocortex

First, we focused on the *H4C3* gene, mutations of which have been reported in a human congenital disorder (TEBIVANED1)(Tessadori et al., 2017), and examined the expression sites of *H4C3* mRNA in mouse embryos. Whole-mount *in situ* hybridization of E10.5 mouse embryos revealed that *H4C3* is expressed throughout the embryo, with particularly strong expression in the limb buds and tail bud (Figure 1A, B). Furthermore, in E14.5 mouse embryos, high levels of *H4C3* expression were observed in highly proliferative tissues, including the spinal cord, liver, phalanges, and brains (Figure. 1C). We also confirmed that other H4C genes, including *H4C1*, *H4C2*, *H4C8*, and *H4C9*, exhibit expression patterns similar to that of H4C3.

**Figure 1.**
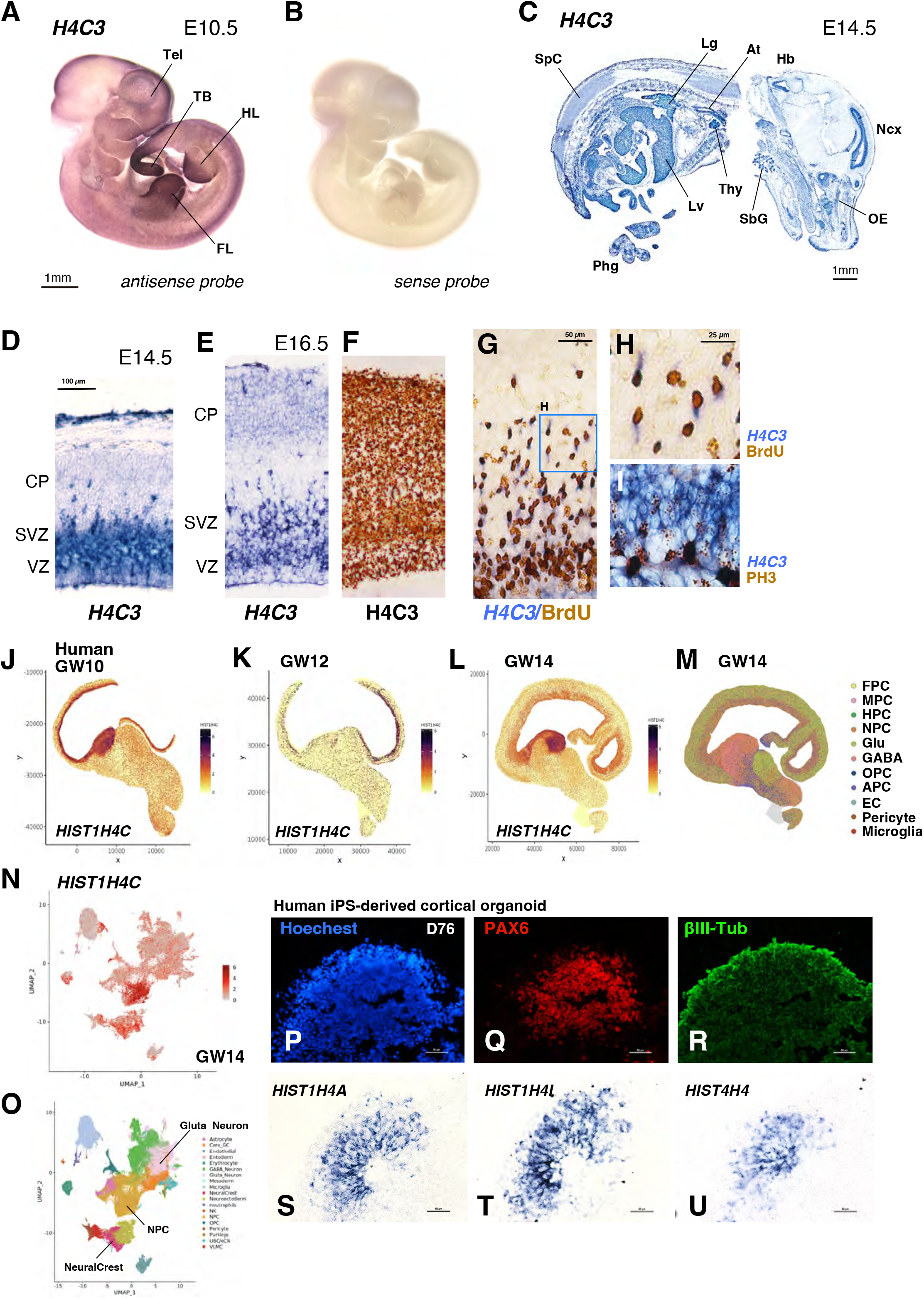
Expression patterns of *H4C* genes in the developing mouse and human brains. (A-C) Expression patterns of *H4C3* mRNA in E10.5 (A, B) and E14.5 mouse embryos. (D-I) Expression patterns of *H4C3* mRNA (D, E) and protein (F) in E14.5 (D) and E16.5 (E) mouse neocortex. (G-I) Co-localization of BrdU or phospho-Histone H3 (PH3) and *H4C3* mRNA in E16.5 mouse neocortex. (J-O) Expression of *HIST1H4C* (J-L) and cell type distribution (M) in Stereo-seq data of the human fetal neocortex. (N, O) Expression of *HIST1H4C* (N) and cell type distribution (O) in sc-RNA data of human fetal neocortex. (P-U) Expression patterns of PAX6, βIII-tubulin (R), *HIST1H4A* (S), *HIST1H4L* (T), and *HIST4H4* (U) in human iPS-derived cortical organoids. Tel: telencephalon; TB: tail bud; FL: forelimb, HL: hindlimb, SpC: spinal cord, Lg: lung; At: atrium; Hb: hindbrain, Lv: liver, Phg: Pha: phalanges; Thy: thymus; SbG: submandibular grand; OE: olfactory epithelium; Ncx: neocortex. CP: cortical plate; SVZ: subventricular zone; VZ: ventricular zone. Scale bars: 1 mm (A, C), 100 µm (D), 50 µm (G), 25 µm (H), 50 µm (P-U).

In the developing neocortex, *H4C3* is highly expressed in the ventricular zone (VZ) and substratum ventricularis (SVZ), where undifferentiated neural progenitor cells are enriched. As development progresses, *H4C3* mRNA was also detected in the cortical plate, which consists primarily of differentiated neurons. The majority of *H4C3*-positive cells in the VZ and SVZ were labeled with BrdU rather than phospho-histone H3, suggesting that *H4C3* is preferentially transcribed during S-phase in proliferating neural progenitors. A public single-cell RNA-seq (scRNA-seq) data revealed that other H4C genes are predominantly expressed in *Pax6-* or *Eomes* (Tbr2)-positive neural progenitors, with weaker expression detected in *βIII-tubulin* (*Tubb3*)-positive neurons, consistent with our *in situ* hybridization analysis (Figure S2). Immunohistochemical analysis using an anti-H4C antibody revealed ubiquitous expression of H4C protein throughout the neocortex, suggesting that H4C protein remain expressed after translation despite the more restricted distribution of their transcripts.

To investigate the expression pattern of H4C genes in the embryonic human brain, we utilized the public database combining spatial transcriptomics and scRNA-seq of the developing human brains. We confirmed that mRNAs of *HIST1H4C* (the ortholog of mouse *H4C3*) is predominantly expressed in the VZ and SVZ of the developing human neocortex (Figure 1J-M). Cell type distribution analysis of the scRNA-seq indicated the higher expression of *HIST1H4C* in neural progenitor cells (NPC) and moderate expression in glutamatergic neurons in GW14 human neocortex (Figure 1N, O). We further corroborate the expression of *HIST1H4* in human iPS-derived cortical organoids. In the organoids on day 76 after neural induction, we confirmed that several *HIST1H4* genes, including *HIST1H4A* (*H1C1*), *HIST1H4I* (*H1C9*), and *HIST4H4* (*H4C16*), were specifically expressed within the neural epithelial structure composed of PAX6-positive cells (Figure 1P-U). These data indicate that the distributions of H4C mRNA in the developing neocortex are evolutionary conserved between rodents and humans.

### 3.2 Disruptions of H4C3 impair skeletal development and neocortical neurogenesis

To further characterize the contribution of *H4C* genes to embryonic development, we generated mice carrying *H4C3* mutations using CRISPR-mediated genome engineering. We designed a gRNA targeting the sequence encoding the C-terminal region of H4C3 and attempted to introduce a non-synonymous substitution by CRISPR-mediated knock-in (Figure S3). However, the resulting animals carried indel mutations that caused frameshifts in the *H4C3* coding sequence and were therefore analyzed as crispants. We then examined skeletal and cerebral cortical development in F0 crispants. Compared with age-matched wild-type embryos, *H4C3* crispants exhibited reduced body size (Figure 2A-G). Skeletal staining revealed that all crispants showed delayed skeletal ossification, and in some cases, cartilage formations in fore- and hind limbs were severely compromised (Figure 2H-M). Immunohistochemical analysis in the developing neocortex revealed an increased abundance of Pax6- and Tbr2-positive progenitor cells and a reduction in βIII-tubulin-positive neurons in the neocortex of *H4C3* crispants (Figure 2N-Q), indicating impaired neuronal production following *H4C3* disruption. These findings suggest that, although H4C transcripts are predominantly expressed in neural progenitor cells, *H4C3* is not essential for the maintenance of radial glial cells and intermediate progenitors but is required for efficient neuronal differentiation from these progenitor populations.

**Figure 2.**
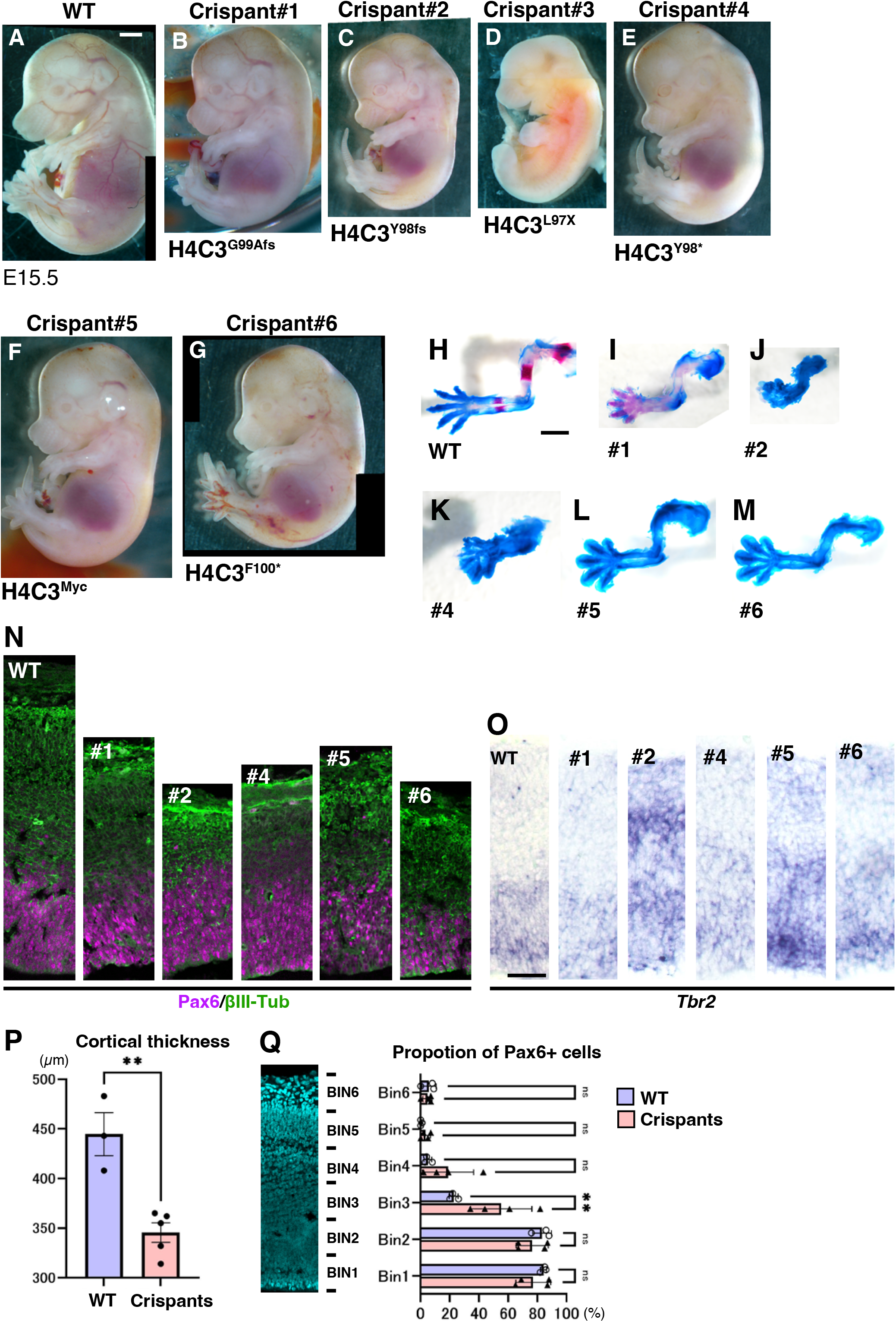
Growth retardation and impaired neurogenesis in H4C3 crispants. (A-G) Gross morphology of E14.5 wild-type (WT; A) and H4C3 crispants (B-G). All crispants were F0 embryos. Since crispant#3 was already dead, it was not possible to conduct further analysis. (H-I) Alucian blue and alizarin red staining of forelimbs of WT and crispants (N, O) Distributions of Pax6, βIII-tubulin (N), and Tbr2 (O) in E14.5 WT and crispants (#1, #2 #4, #5, #6) neocortex. (P, Q) Cortical thickness (P) and distributions of Pax6-positive cells (Q) in WT and crispant neocortex. WT: n=3 samples; cirspants: n=5 (#1, #2 #4, #5, #6).

### 3.3 A patient-type H4C3 mutation mis-regulates genes associated with epithelial cell development in the developing brains

Previous studies suggest that a non-synonymous amino acid substitution (K91Q or K91R) in H4C3 associated with TEBIVANED1 syndrome results in various neurodevelopmental abnormalities (Tessadori et al., 2017). To investigate the effect of the patient-type H4C mutation, we examined the effects of H4C3K91Q in the regulation of downstream gene expression in the developing mouse brains. GAGE (Generally Applicable Gene-set Enrichment) analysis of a published RNA-seq data of iPS-derived neurons carrying H4C3K91Q (Feng et al., 2023) revealed that a patient-type H4C3 mutation downregulates genes associated with specific biological terms, including “positive regulation of cell motility” or “positive regulation of cell migration” (Figure 3A). To confirm whether the H4C3 mutation affects cellular characteristics during brain development, we overexpressed the vectors expresses H4C3K91Q into the E13.5 mouse neocortex by *in utero* electroporation. The distribution of GFP-positive cells in the developing neocortex were not significantly changed by K91Q mutant H4C in comparison with the control embryos with GFP-expression vector alone (Figure 3B-E). However, we identified that H4K91Q induced the clump of GFP-positive cells in the intermediate zone of the neocortex (n=2 out of 3 embryos; Figure 3C), which were not observed in control embryos (n=3 embryos; Figure 3B). Furthermore, compared to controls, K91Q overexpression resulted in GFP accumulation in the basal endofeet of radial glial cells (RGCs; Figure 3F, G), suggesting enhancement of polarized characteristics of RGCs. To identify the genes directly affected by the mutant H4C in the developing neocortex, we performed bulk RNA-seq analysis of neocortical cells transfected with wild-type H4C3 (H4C3WT) and H4C3K91Q (Figure 3H). We identified total 26 genes that were up- or down-regulated by H4C3K91Q (Figure 3I and Table S2).

**Figure 3.**
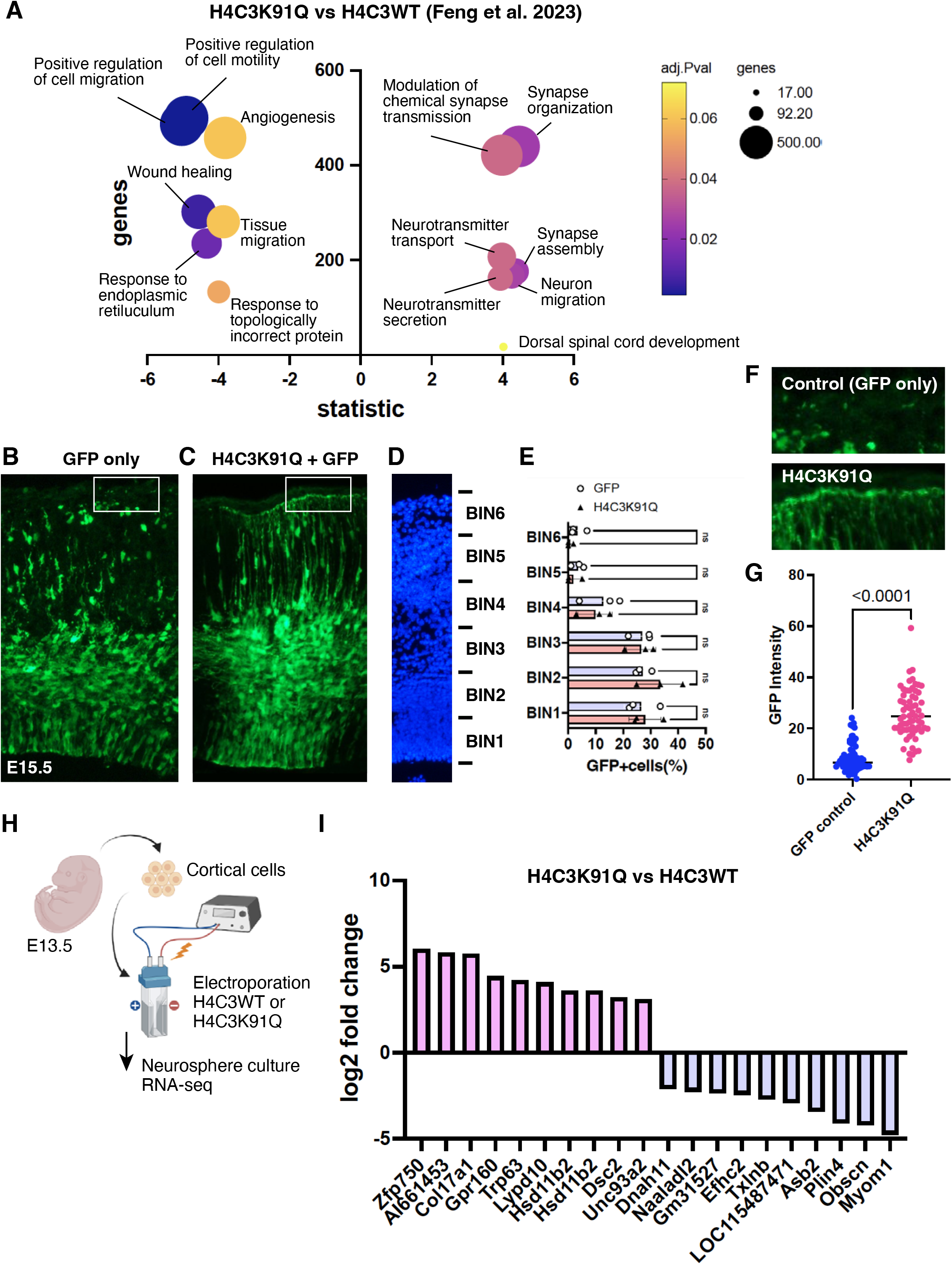
Altered gene expression by H4C3K91Q mutant. (A) GAGE analysis of bulk RNA-seq data by Feng et al (Feng et al. 2023). (B, C) Overexpression of GFP (B) and H4C3K91Q (C) expression vectors into the developing neocortex. *In utero* electroporation was performed at E13.5, and embryos were collected at E15.5. Arrowhead indicate Ectopic cell aggregations. (D, E) Distributions of GFP-positive cells in embryos overexpressed with control and H4C3K91Q expression vectors. (F) GFP distributions in the endfeets of control and H4C3K91Q-intoduced neocortex. (G) Quantification of GFP intensity of endfeets in control and H4C3K91Q-intoduced neocortex. (H, I) bulk RNA-seq analysis of mouse neocortical cells transfected with H4C3WT and H4C3K91Q mutant. (H) After electroporation with WT or mutant histone H4Cells were cultured in neurosphere medium for 1 day, and performed RNA-seq analysis. (I) Up or down-regulated genes by H4C3K91Q mutant compared with H4C3WT. Top 10 genes with log2 FC >|2|.

Among these, several genes including *Zfp750*, *Coll17a1*, *Hsd11b2*, and *Myom1* were overlapped with genes identified in a previous transcriptome analysis (Table S2). Notably, *Zfp750* and *Coll17a1*, which are responsible for the development of epidermal cells and basement membranes (Liu et al., 2024; Tanimura et al., 2011). These data suggest that a patient-type H4C mutation mis-regulates the expression of genes associated with epithelial cell development and/or maintenance, in the developing neocortical cells.

## 4. Discussion

Here we confirmed unique expression patterns of H4C genes in the developing mouse and human brains. H4C genes are highly transcribed in the neural progenitors reside in the VZ and SVZ, and also weakly expressed in differentiated neurons. These expression patterns are contrast to the ubiquitous distributions of H4C protein in the developing neocortex, suggesting that transcription and translation of H4C genes are tightly controlled by cell type-specific regulatory mechanisms.

The molecular mechanisms underlying the S-phase-specific expression of histone genes have been extensively characterized (Banday et al., 2014; Braastad et al., 2004; Marzluff et al., 2008). Histone gene transcription is coordinated by histone locus bodies (HLBs), which are organized around nuclear protein ataxia-telangiectasia locus (NPAT) (Ye et al., 2003). At the G1/S transition, the cyclin E–CDK2 complex phosphorylates NPAT, triggering the recruitment of multiple transcriptional regulators, including histone nuclear factor P (HINFP) (Ghule et al., 2018; Miele et al., 2005). HINFP binds to the Site II element in histone gene promoters and recruit RNA polymerase II, thereby activating histone transcription. Phosphorylated NPAT has also been proposed to recruit the Tip60/TRRAP histone acetyltransferase complex, resulting in chromatin remodeling and enhanced transcriptional activation of histone genes (DeRan et al., 2008). In addition, the histone acetyltransferase 1 (HAT1) holoenzyme associates with active H4 gene promoters. HAT1 is thought to couple cellular metabolic status to histone gene expression by utilizing acetyl-CoA generated from glucose metabolism, thereby linking nutrient availability to the initiation of histone H4 transcription (Gruber et al., 2019). Following transcription, multiple HLB-associated proteins and RNA-processing factors regulate histone mRNA maturation, nuclear export, stability, and translation. At the completion of S phase, excess replication-dependent histone mRNAs are rapidly degraded through a pathway involving the histone-specific 3ʹ–5ʹ exonuclease (3ʹhExo), thereby preventing inappropriate histone protein synthesis outside S phase (Meaux et al., 2018).

Replication-dependent histone genes lack introns, and their transcripts are not polyadenylated. Instead, the 3ʹ untranslated region contains a highly conserved stem-loop structure that is recognized by the stem-loop-binding protein (SLBP), which promotes the nuclear export, stability, and efficient translation of mature histone mRNAs, enabling the rapid synthesis of histone proteins during S phase (Marzluff et al., 2008). On the other hand, in post-mitotic cells such as differentiated neurons, the replication-dependent histone genes *H4C8* and *H4C14/15* have been shown to maintain histone H4 protein expression independently of the cell cycle by producing polyadenylated transcripts (Lyons et al., 2016). Therefore, it is possible that the H4C mRNA detected in the cortical plate of the developing mouse cerebral cortex represents a polyadenylated transcript rather than the canonical non-polyadenylated replication-dependent histone mRNA.

We demonstrate that disruption of H4C3 in mice editing results in severe developmental abnormalities, including growth retardation, delayed skeletal ossification, and microcephaly, which partially resemble to the zebrafish or mice with human patient-type H4C mutations. In the H4C3 crispants, Pax6 and Tbr2-positive neural progenitors are maintained, while differentiation neurons were severely reduced, suggesting that H4C3 plays critical roles in neuronal differentiation, rather than progenitor proliferation. These phenotypes contrast to the precocious neuronal differentiation by H4K19Q or H4K91R mutations (Feng et al., 2023), suggesting that distinct developmental pathways are affected by the patient-type H4C mutations. Notably, we identified that H4C3K91Q significantly up-regulates specific category of genes, such as *Zfp750* and *Coll17a1*, which are associated with epidermal development and skin cell adhesion. Concomitantly, overexpression of H4C3K91Q induced abnormal neuronal clusters and enhanced basement attachments of radial glial cells. It has been shown that H4C3K91 mutant protein is preferentially associated with H3.3 in DAAX/ATRX-dependent manner, and increased chromatin accessibilities. Collectively, these lines of evidence suggest that incorporation of patient-type H4C mutants into the nucleosome complex results in aberrant gene expression in neuronal progenitors, by which consequences disturbance of cell fates in the developing neocortex. A recent study reported that in the developing human brains, all H4C genes are initially expressed in almost equivalent proportion, while *H4C4* (*HIST1H4D*), *H4C8* (*HIST1H4H*), and *H4C16* (*HIST4H4*) are predominantly transcribed as development proceed (Matheson-Grant, 2024). Thus, the expression of each H4C genes are also tightly regulated in a stage-specific manner. Further investigation of spatio-temporal regulatory mechanisms of H4C genes and their downstream cascades are necessary to understand the developmental roles of H4C genes in normal and pathogenic conditions.

## Statement

### Data availability statement

The original contributions of presented in the study are included in the article/Supplementary Material, further inquiries can be directed to the corresponding author. The raw data of RNA-seq has been deposited to DDBJ (PRJDB42636).

### Ethics statement

All animal experiments were approved by the experimental animal committee of Kyoto Institute of Technology (#100211) and were performed in accordance with the relevant guidelines.

### Author contributions

HN: Investigation, Formal Analysis, Writing-original draft. KN: Methodology, Writing-review and editing, ST: Writing-review and editing, TN: Writing-original draft, review and editing, Formal Analysis, Project Administration, Conceptualization, Funding acquisition, Supervision.

## Funding

The author(s) declared that financial support was received for this work and/or its publication. This work was supported by Terumo Life Science Foundation, Takeda Science Foundation, KIT cutting-edge science project, and Integrative Center for Kyoto Health Science 2024, 2025 and 2026. The funders has no role in study design, data collection, and analysis, decision to publish, or preparation of the manuscript.

## Acknowledgments

We thank Ms. Misato Kawami, Mariko Yazaki and Akiko Watanabe for technical assistance, and Dr. Ryota Noji for advising RNA-seq analysis. We also appreciate all members of Biomedical and Developmental Biology lab for helpful discussion on the work. A part of Figures are constructed by BioRender [YD29VBFB4, Nomura, T. (2026) https://BioRender.com/zq1warf]

## Conflicts of interests

The author(s) declared that this work was conducted in the absence of any commercial of financial relationships that could be constructed as a potential conflict of interest.

## Generative AI statement

The author(s) declared that generative AI was not used in the creation of the manuscript.

## Notes

### Competing Interest Statement

The authors have declared no competing interest.

