## Supplementary Figures for "Disruption of *Histone H4C* genes impairs skeletal development and cortical neurogenesis, modeling rare neurodevelopmental syndromes"

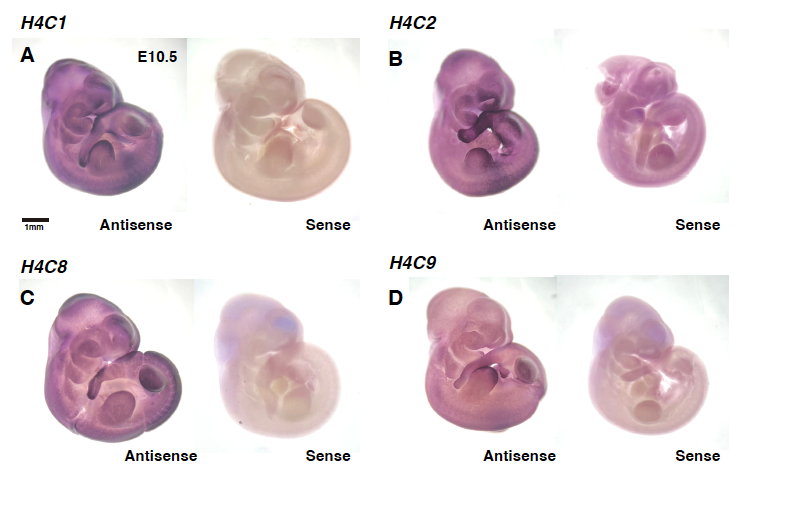

**Figure S1. *H4C* genes are highly expressed in early embryogenesis**

Whole mount *in situ* hybridization of E10.5 mouse embryos with *H4C1* (A), *H4C2* (B), *H4C8* (C), and H4C9 (D) probes. Scale bar: 1 mm (A).

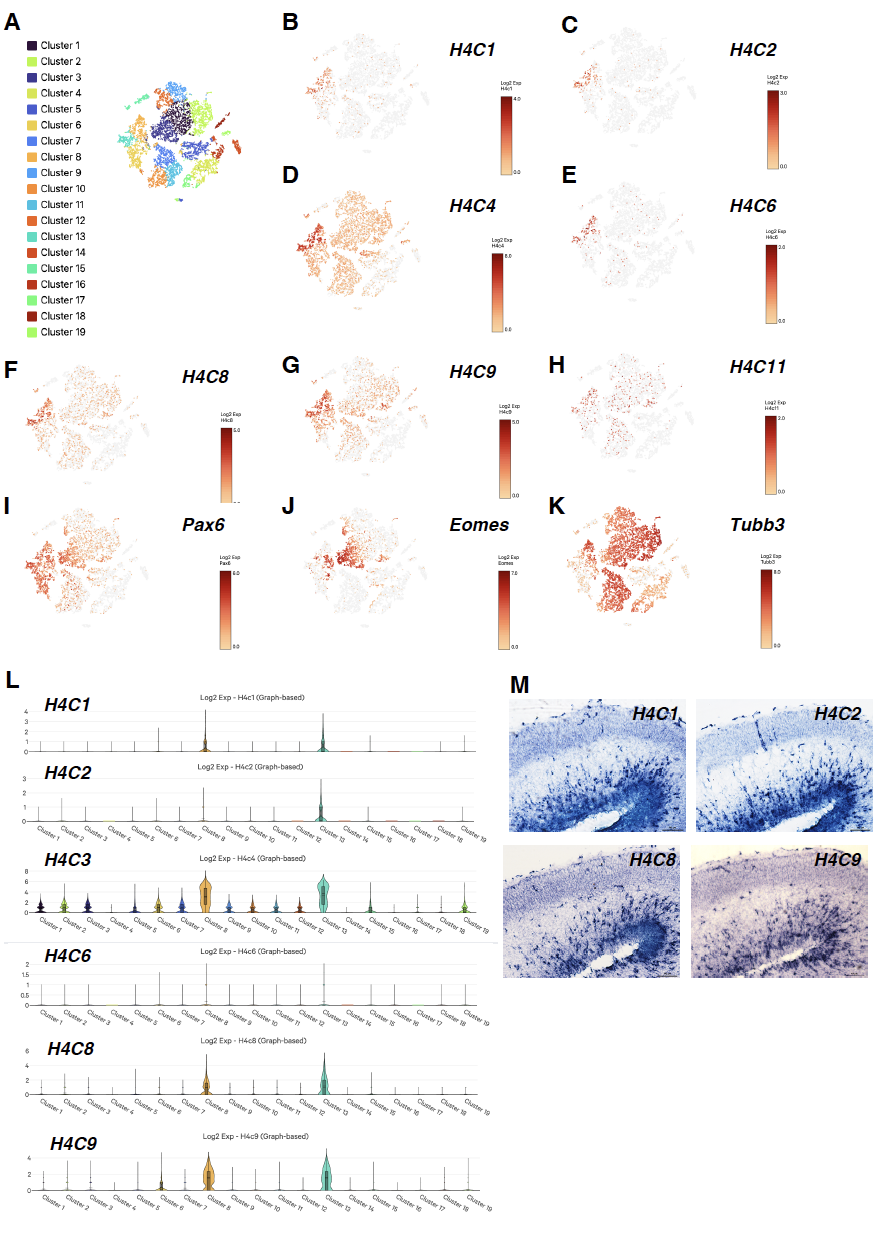

**Figure S2. *H4C* genes are highly expressed in mouse cortical neural progenitors**

(A-L) scRNA-seq data of E18 mouse neocortex (10xGenomics). Cell type distributions with different clusters are shown in (A). The expression of H4C genes are shown in (B-H, L). *Pax6* (I), *Eomes* (J) and *Tubb3* (K) represent radial glial cells, intermediate progenitors, and differentiated neurons, respectively. (M) *In situ* hybridization of *H4C1*, *H4C2*, *H4C8*, and *H4C9* in E16.5 mouse neocortex.

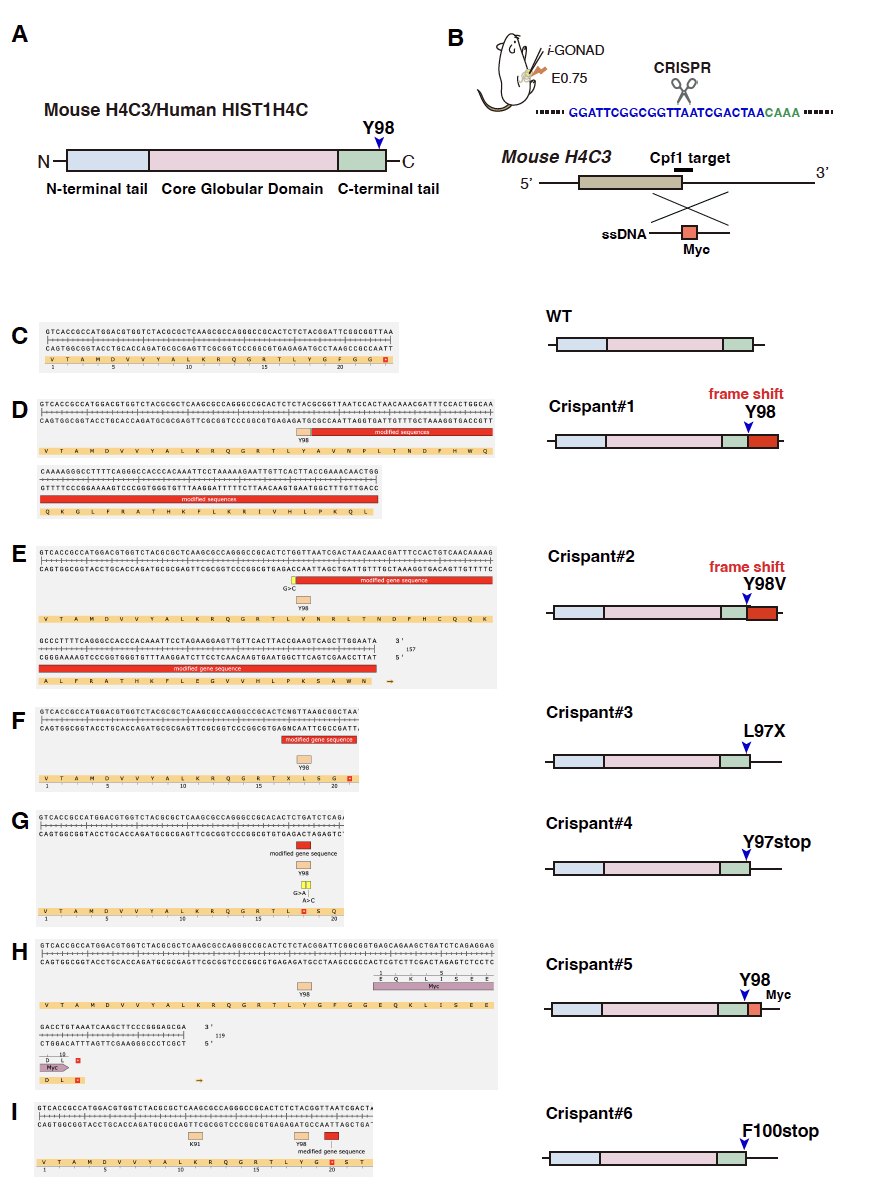

**Figure S3. Frame shift mutations in *H4C3* crispant embryos**

(A) Protein structure of mouse H4C3 and human *HIST1H4C*. H4C consists of N-terminal tail, core globular domain, and C-terminal tail. (B) A target sequence of gRNA in mouse H4C3 gene. Crispr/Cpf1-mediated genome editing was performed into fertilized eggs by using *i*-GONAD method. Myc-tagged ssDNA was introduced to induce homologous recombination. (C-I) Sequences of H4C3 C-terminal tails in WT (C) and crispants (D-I).
