## Supplementary Table 1 for "Disruption of *Histone H4C* genes impairs skeletal development and cortical neurogenesis, modeling rare neurodevelopmental syndromes"

**Table S1. Sequences of primers and gRNA for CRISPR-mediated gene targeting**

| **Name of primers/crRNA** | **Sequences** |
| --- | --- |
| H4C1 forward (mouse) | 5’-CGGTATCGATAAGCTTGTGATTCTGTTGGG-3’ |
| H4C1 reverse (mouse) | 5’-ATTCGATATCAAGCTTGGGCGGCCCTGAGA-3’ |
| H4C2 forward (mouse) | 5’-CGGTATCGATAAGCTCAACCAGGTCCGATA -3’ |
| H4C2 reverse (mouse) | 5’-ATTCGATATCAAGCTTGAGTGGTCCTGAAA -3’ |
| H4C8 forward (mouse) | 5’-CGGTATCGATAAGCTGAAAACACTGCATAT-3’ |
| H4C8 reverse (mouse) | 5’-ATTCGATATCAAGCTTGGGCGGCCCTGAAA-3’ |
| H4C9 forward (mouse) | 5’-CGGTATCGATAAGCTATCTGCTTCGTCTTA-3’ |
| H4C9 reverse (mouse) | 5’-ATTCGATATCAAGCTTGTGGTGGCCCTAAA-3’ |
| HIST1H4A forward (human) | 5’-CGGTATCGATAAGCTACTTGCTCTTGGTTC-3’ |
| HIST1H4A reverse (human) | 5’-ATTCGATATCAAGCTTGGGCGGCCCTGAAA-3’ |
| HIST1H4C forward (human) | 5’-CGGTATCGATAAGCTACTGCGATAGGAATC-3’ |
| HIST1H4C reverse (human) | 5’-CGGTATCGATAAGCTTGAGCGGCCCTGAAA-3’ |
| HIST1H4I forward (human) | 5’-CGGTATCGATAAGCTAGACCTTTGTTCTCT-3’ |
| HIST1H4I reverse (human) | 5’-ATTCGATATCAAGCTAAGCTTAGGGGCCCT-3’ |
| HIST4H4 forward (human) | 5’-CGGTATCGATAAGCTACAGTGGGCAGCCCG-3’ |
| HIST4H4 reverse (human) | 5’-ATTCGATATCAAGCTTGGGGGCCCTGAAAA-3’ |
| H4C3 crRNA | 5'-[TTTG(PAM)]TTAGTCGATTAACCGCCGAATCCG-3 |
